## Supplemental_Figures for "*In situ* dissection of domain boundaries affect genome topology and gene transcription in *Drosophila*"

### Supplementary Figures 1-6

#### Supplementary Fig. 1

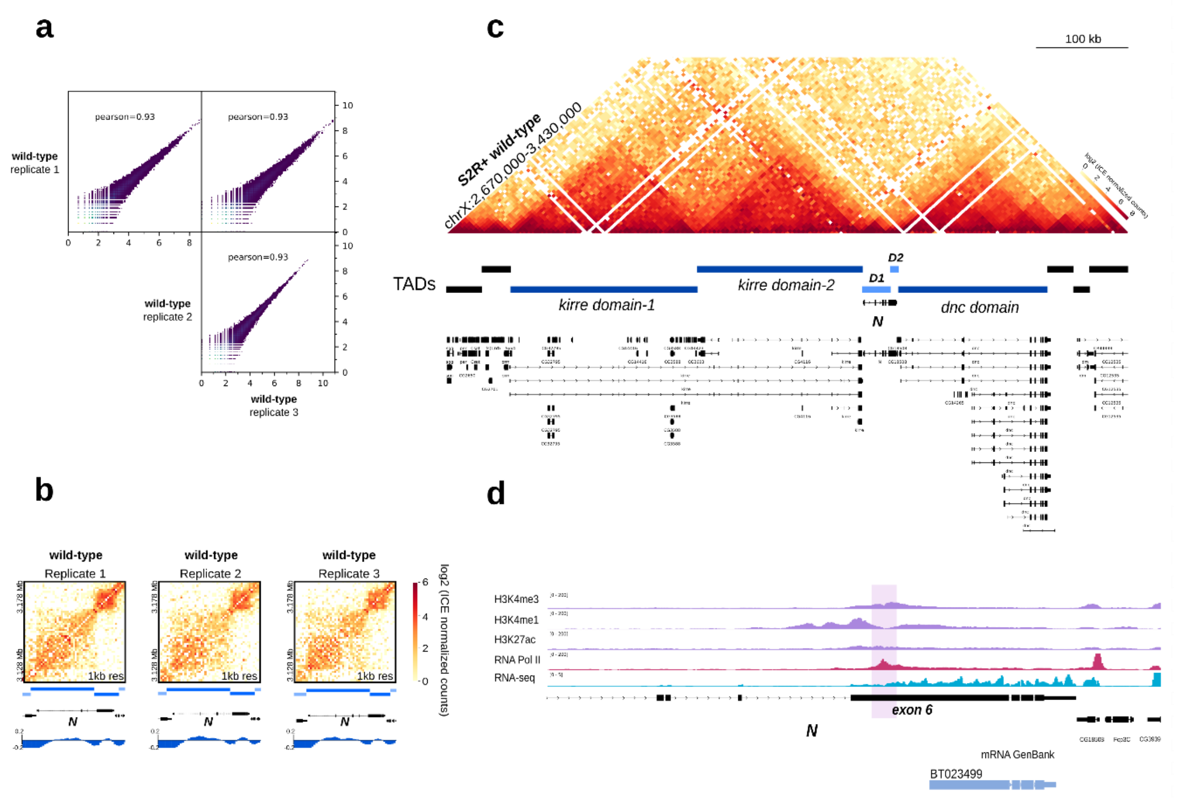


**Supplementary Fig. 1. Topological domains spanning the *kirre* and *dnc* loci flank *Notch***

**a** Scatterplots comparing interactions between wild-type Hi-C replicates. The Pearson correlation value for each pair-wise comparison is shown.

**b.** Hi-C normalized heatmaps at 1 kb resolution covering a 50 kb region centered in *Notch* for each wild-type replicate. The position of TADs and the TAD separation score for each dataset are shown below each heatmap.

**c** Hi-C heatmap at 5 kb resolution showing the topological landscape surrounding *Notch*. TADs identified at 1kb resolution are shown below the heatmap. The *kirre* locus is partitioned into two TADs termed *kirre* domain-1 and 2 while the *dnc* locus is fully contained within a TAD.

**d** Genome browser track displaying part of the *Notch* locus with public ChIP-seq data for histone marks, RNA Pol II and RNA-seq in S2R+ cells. The putative exonic promoter is highlighted. Below a cDNA identified for this region and reported in GenBank is shown.

### Supplementary Fig. 2

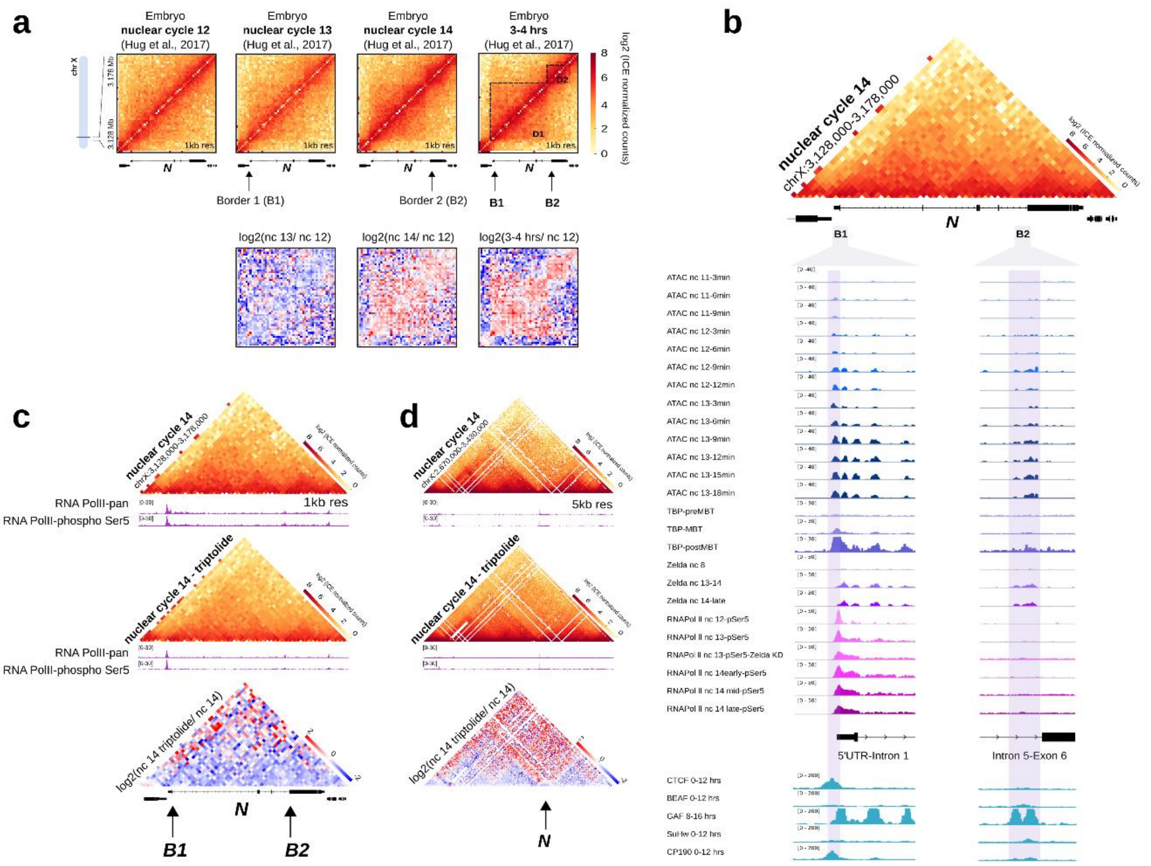


**Supplementary Fig. 2. *Notch* 3D organization emerges during early embryonic development and correlates with the gain of chromatin accessibility and binding of RNA Pol II at domain boundaries**

**a** Hi-C heatmaps at 1 kb resolution covering a 50 kb region centered in *Notch*. Hi-C from developing embryos (nuclear cycle 12,13,14 and 3-4 hrs of development; Hug et al., 2017) was re-analyzed using the same pipeline used to analyze the Hi-C data generated in this study. Black arrows indicate the position of B1 and B2 boundaries of *Notch.* Dotted lines indicate the position of the D1 and D2 domains of *Notch*. Below, Hi-C heatmaps of the log2 differences in interaction frequency between the nuclear cycle 12 (nc 12) and nc13, nc14, and 3-4 hrs.

**b** Triangular representation of a Hi-C heatmap from nuclear cycle 14 wild-type embryos at 1 kb resolution covering a 50 kb region centered in *Notch.* Below the heatmap are shown tracks for public ATAC-seq, and ChIP-seq datasets for TBP, RNA Pol II, and Zelda at different time points during early embryonic development (nc11-nc14) for the regions identified as boundaries at the *Notch* locus. ChIP-seq data for Architectural Proteins from 0-12 hrs embryos is also shown. MBT, mid blastula transition. Note that the B1 domain boundary is highly accessible and shows a strong enrichment or RNA pol II and TBP.

**c** Triangular representation of Hi-C heatmaps at 1 kb resolution covering a 50 kb region centered in *Notch*. *Top* and *middle*, Hi-C from nuclear cycle 14 wild-type or triptolide treated embryos. Below each heatmap are shown RNA Pol II ChIP-seq tracks from wild-type, and triptolide treated embryos from Hug et al., 2017. *Bottom*, Hi-C heatmaps of the log2 difference in interaction frequency between wild-type nuclear cycle 14 and nuclear cycle 14 triptolide treated embryos. Black arrows indicate the position of the B1 and B2 boundaries of *Notch.* Observe that global transcriptional inhibition results in a decrease of intradomain interactions at the *Notch* locus; however, TADs are still visible.

**d** Triangular representation of Hi-C heatmaps at 5 kb resolution covering a 760 kb region centered in *Notch. Top* and *middle*, Hi-C from nuclear cycle 14 wild-type or triptolide treated embryos. Below each heatmap are shown RNA Pol II ChIP-seq tracks from wild-type, and triptolide treated embryos from Hug et al., 2017. *Bottom*, Hi-C heatmaps of the log2 difference in interaction frequency between wild-type nuclear cycle 14 and nuclear cycle 14 triptolide treated embryos. Black arrow indicates the position of *Notch.* Observe that global transcriptional inhibition results in a decrease of intradomain interactions also at TADs flanking the *Notch* locus; however, TADs are still visible.

### Supplementary Fig. 3

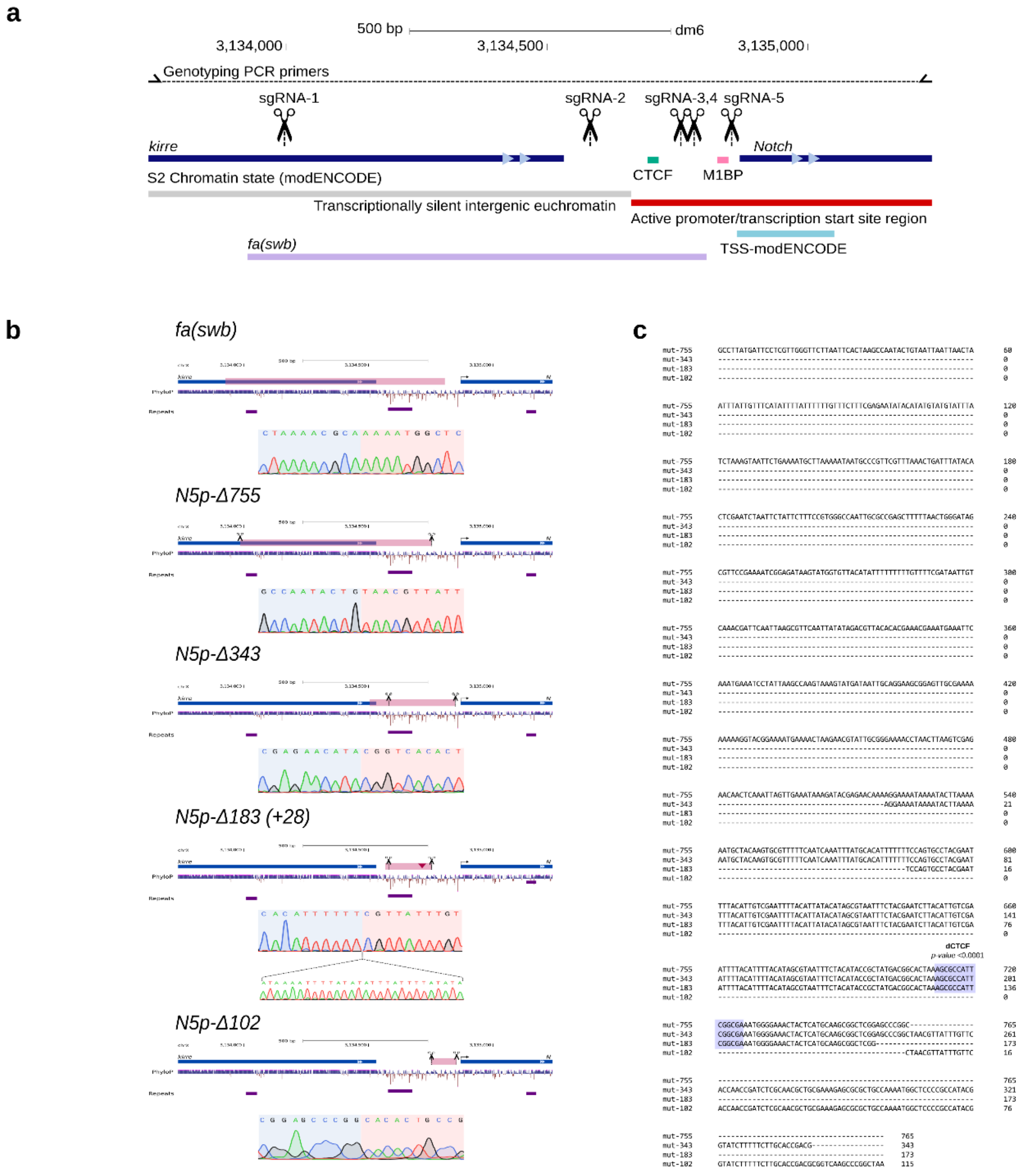


**Supplementary Fig. 3. CRISPR-Cas9 strategy for deletion of the B1 boundary and sequencing results for mutant clones**

**a** Design for CRISPR/Cas9 mediated deletions over the 5’ intergenic region of *Notch*.

**b** Sanger-sequencing breakpoints for all mutant alleles generated by CRISPR-Cas9 in S2R+ cells as well as the breakpoints of the *fa(swb)* mutant flies. Pink boxes represent the deleted sequence in each mutant. Scissors represent the sgRNAs used to generate each deletion. For each mutant an electropherogram of the sequencing results at the deletion breakpoints is shown. A red triangle represents an insertion for the Δ183 mutant allele.

**c** Multiple alignment of the deleted sequences from each 5' mutant generated in this study. Note that with exception of the 102 allele, all other mutant alleles loss the binding site for CTCF.

### Supplementary Fig. 4

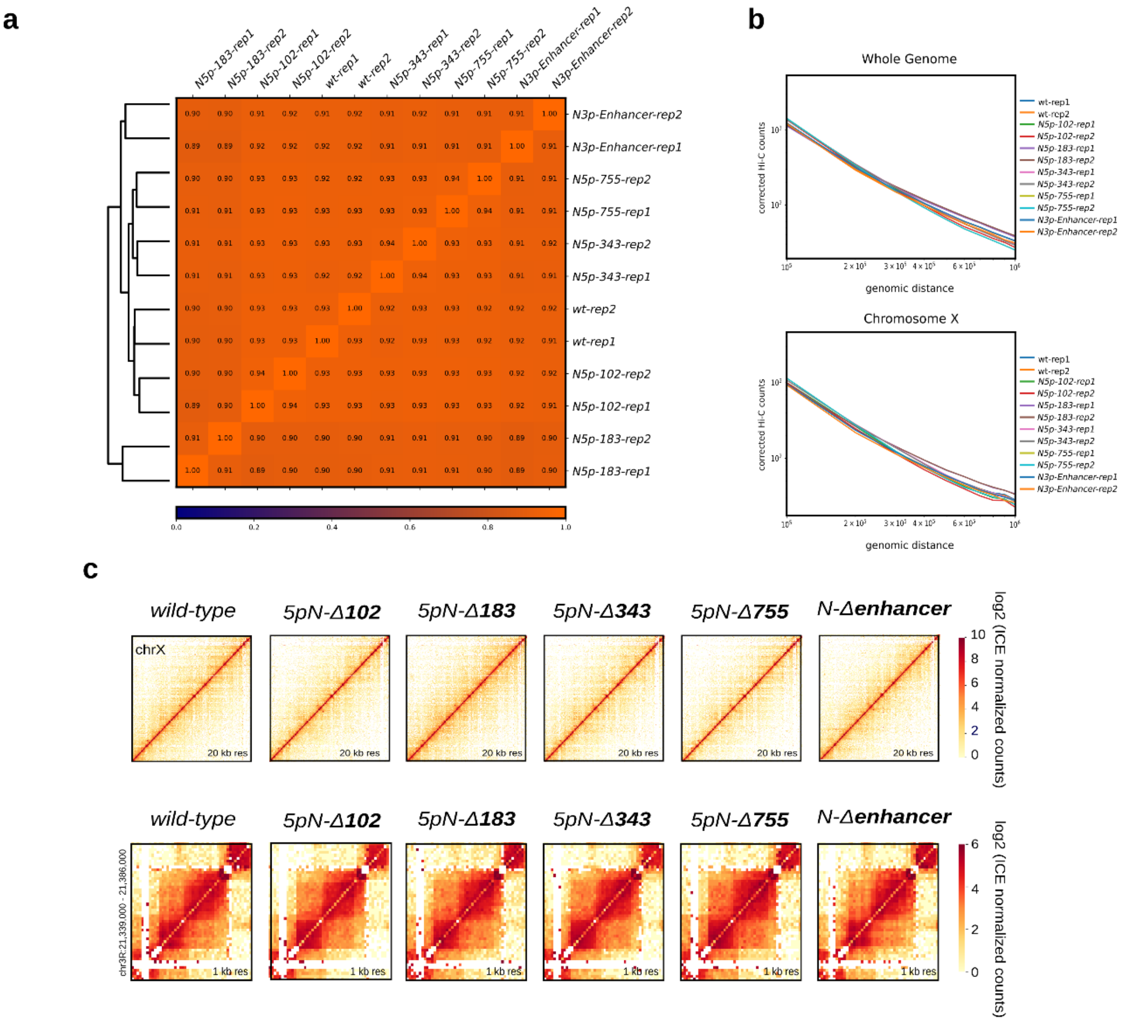


**Supplementary Fig. 4. Quality Control for Hi-C data from CRISPR mutant clones**

**a** Pearson correlation heatmap between all Hi-C datasets generated in this study.

**b** Distance vs HiC counts plots for all Hi-C data sets generated in this study. *Top*, whole-genome. *Bottom*, X chromosome.

**c** *Top*, Hi-C heatmaps at 20kb resolution for the X chromosome for wild-type and all CRISPR mutants generated in this study. *Bottom*, Hi-C heatmaps at 1 kb resolution centered in *mod(mdg4)* for wild-type and all CRISPR mutants. Observe that *mod(mdg4)* is organized into a TAD with additional subTADs spanning the locus. CRISPR deletion at *Notch* domain boundaries on the X chromosome do not affect the overall organization of the X chromosome nor the organization of a locus in a different chromosome.

### Supplementary Fig. 5

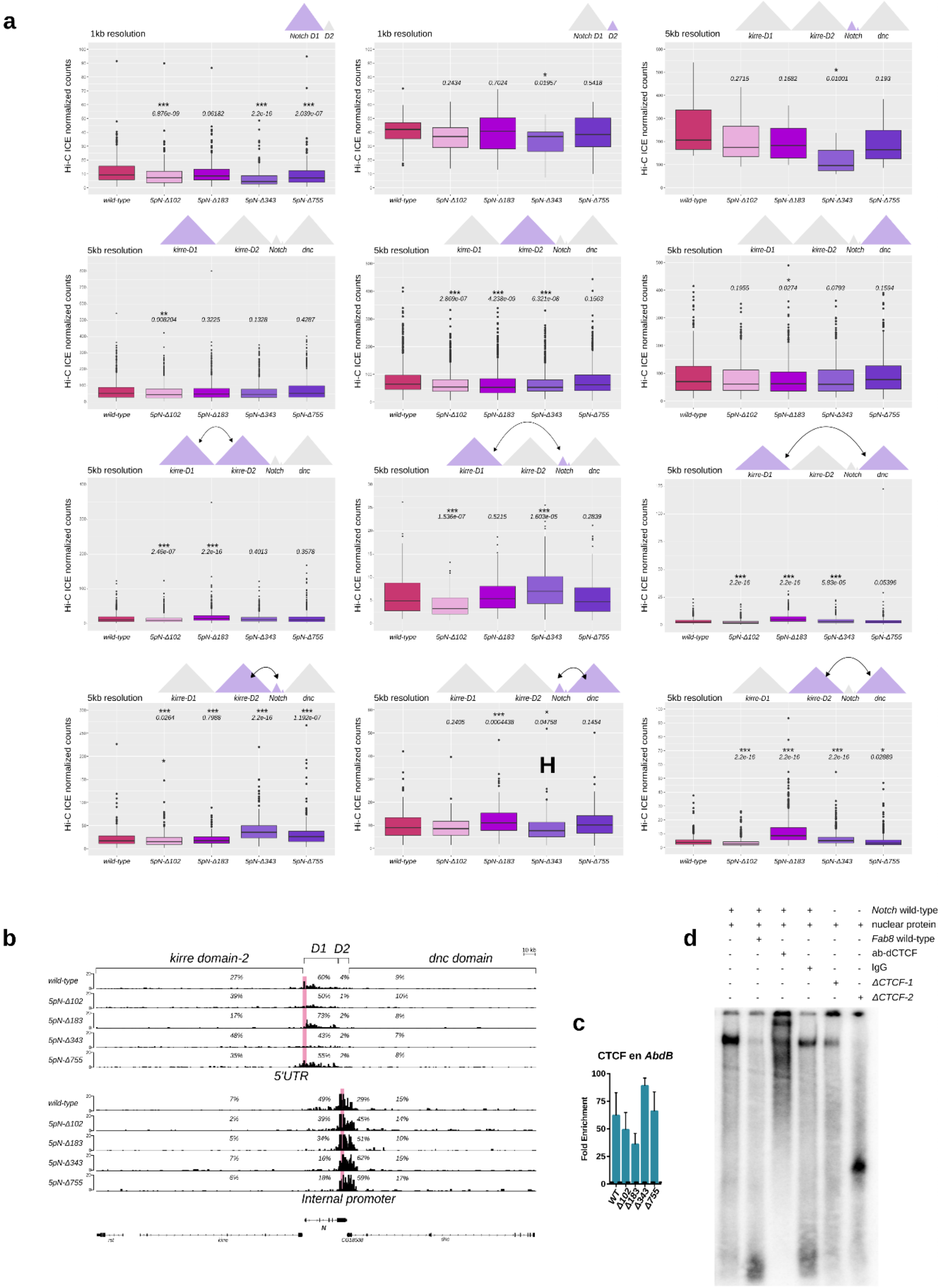

**Supplementary Fig. 5. Deletion of the B1 boundary of *Notch* results in topological defects**

**a** Boxplots of Hi-C normalized counts for *kirre* domain-1 and 2, *Notch* domains and *dnc* domain as well as inter-domain interactions, in wild-type and B1 boundary CRISPR mutants. On top of each plot a diagram depicting the location of the evaluated interactions is shown. *p-values* from a Wilcoxon-Rank Sum Test are shown on top of each boxplot. *p-value* *<0.05, **<0.01,***<0.001.

**b** Virtual 4C of Hi-C data using the 5'UTR and the Internal Promoter of *Notch* as viewpoints for the wild-type and B1 boundary CRISPR mutants. Percentages in each track indicates de fraction of valid-reads interacting with the viewpoint in each domain.

**c** qChIP against CTCF using a set of primers for the control region *AbdB* for wild-type and CRISSPR mutants. Shown are fold enrichment values over IgG.

**d** EMSA using S2R+ protein nuclear extracts and oligonucleotides listed in Table 2. As a control, a non-labeled 60 bp oligonucleotide with a well-validated CTCF binding motif from the *Fab 8* insulator was used. From left to right; 1, shift with the *Notch* wild-type oligonucleotide; 2, competition of the labeled *Notch* wild-type oligonucleotide with the non-labeled *Fab8* oligonucleotide; 3, super-shift assay with protein nuclear extracts incubated with α-dCTCF and the *Notch* wild-type oligonucleotide; 4, incubation of wild-type oligonucleotide with IgG; 5 and 6 CTCF mutant oligonucleotides incubated with protein nuclear extracts. Note that mutant oligonucleotides either reduce the binding of nuclear proteins (ΔCTCF-1) or completely disrupt wild-type shift (ΔCTCF-2).

### Supplementary Fig. 6

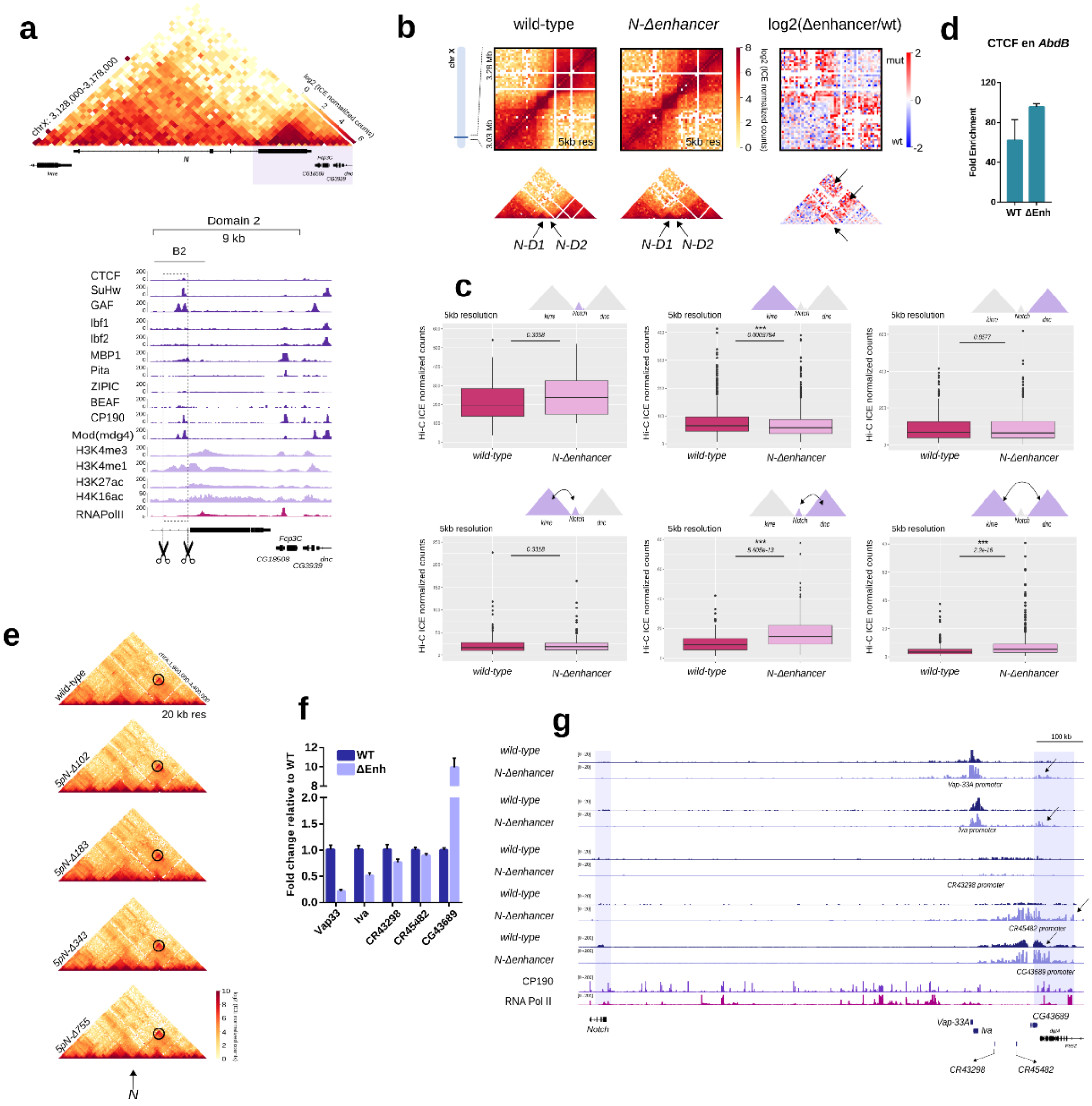


**Supplementary Fig. 6. Deletion of the B2 boundary of *Notch* results in local and long-range topological defects.**

**a** The B2 boundary of *Notch* overlaps an intronic enhancer occupied by CP190 and other Architectural Proteins. A triangular representation of a Hi-C heatmap at 1 kb resolution covering a 50 kb region and centered in *Notch* is shown on top. Highlighted is a region covering from the B2 boundary to the end of the D2 domain of *Notch*. This genomic region is shown below with ChIP-seq tracks for APs, histone marks, and RNA Pol II. Scissors represent the position of sgRNAs used for CRISPR-Cas9 mediated deletion of the intronic enhancer.

**b** Deletion of the intronic enhancer results in increased inter-domain interactions between TADs flanking *Notch*. *Left* and *center*, heatmaps at 5 kb resolution of Hi-C data for wild-type and the enhancer mutant covering a region of ~250 kb centered in *Notch*. *Right*, heatmap of the log2 differences between wild-type and the enhancer mutant. The arrows in highlight regions that show a gain of interactions in the enhancer mutant.

**c** Boxplots of Hi-C normalized counts for *kirre*-domain 1 and 2, *Notch* domains and *dnc*-domain as well as inter-domain interactions in wild-type and the enhancer mutants. On top of each plot, a diagram depicting the location of the evaluated interactions is shown. *p-values* from a Wilcoxon-Rank Sum Test are shown on top of each boxplot. *p-value* *<0.05, **<0.01,***<0.001.

**d** qChIP against CTCF using a set of primers for the control region AbdB for wild-type and enhancer mutants. Fold enrichment over IgG.

**e** Triangular representation of Hi-C heatmaps covering a 2.5 Mb region at 20 kb resolution for wild-type cells and the B1 boundary CRISPR mutants centered in *Notch*. Highlighted by a black circle is a 1 Mb long-range interaction mediated by the intronic enhancer of *Notch*.

**f** Transcription of genes located at or near the gene desert interacting with the intronic enhancer of *Notch* in wild-type and mutant cells. Significant differences between wild-type and enhancer mutant were calculated using a t-test. n=3, *p-value* *<0.05, **<0.01,***<0.001.

**g** Virtual-4C for wild-type and the enhancer mutant using as viewpoints the promoter of genes located at or near the gene desert interacting with the intronic enhancer of *Notch* in wild-type and mutant cells.  Arrows indicate regions with ectopic interactions. ChIP-seq tracks for CP190 and RNA Pol II are shown.
